# Challenges in establishing epidemiological cut-off values for the *Burkholderia cepacia* complex

**DOI:** 10.64898/2026.05.18.725987

**Authors:** Holly K. Huse, Carmila Manuel, Tracy McLemore, Romney M. Humphries, Anna Clara Milesi Galdino, Diana Celedonio, John J. LiPuma, Daniel A. Green, James E. A. Zlosnik, Maria Traczewski, Audrey Schuetz, John Turnidge, Mandy Wootton, Darcie Carpenter, Michael Huband, Chris Pillar, Marguerite L. Monogue, Peter A. Jorth

## Abstract

The *Burkholderia cepacia* complex (BCC) is comprised of 24 species of Gram-negative bacteria that cause opportunistic infections. While antimicrobial susceptibility testing (AST) has historically been used to guide treatment for BCC infections, recent work highlighting problems with AST for these organisms led the Clinical and Laboratory Sciences Institute (CLSI) to remove disk diffusion (DD) and minimal inhibitory concentration (MIC) breakpoints for BCC from its M100 standards document. Epidemiological cut-off values (ECVs) may be helpful to clinicians in the absence of breakpoints, as they may be used to determine whether an isolate has a wild-type or non-wild-type phenotype. Here we present an analysis of BCC ECVs for ceftazidime (CAZ), levofloxacin (LVX), meropenem (MEM), minocycline (MIN), and trimethoprim-sulfamethoxazole (TMP-SMX). ECVs were calculated using MIC data from 3 previous studies and 3 independent laboratories for 1,896 BCC isolates. ECVs were 16 μg/ml for CAZ, 8 μg/ml for LVX, 16 μg/ml for MEM, and 8 μg/ml for MIN. The ECV for TMP-SMX varied depending on the analysis from 2 μg/ml, 8 μg/ml, and 16 μg/ml and therefore could not be reliably established. Challenges with establishing ECVs for BCC include limitations with the pooled MIC dataset, broad MIC distributions, and high ECVs that are above the obsolete susceptible MIC breakpoints. These challenges limit the clinical utility of ECVs for these organisms and supported removal of ECVs from the CLSI M100 standards document.

**IMPORTANCE:** The *Burkholderia cepacia* complex is a group of bacterial species that cause difficult-to-treat opportunistic infections. Recently, clinical breakpoints, which are used to determine whether organisms are susceptible to certain antimicrobials, were removed from Clinical and Laboratory Standards Institute (CLSI) standards for these organisms due to problems with antimicrobial susceptibility testing performance. Clinicians are now faced with the challenge of how to treat these complex infections without clinical breakpoints. Here we determine epidemiological cut-off values (ECVs) for relevant antimicrobials for the *B. cepacia* complex. While we established ECVs for four antimicrobials, we encountered significant challenges in our analyses, including limitations with data for these organisms and high ECVs that are not clinically useful. These challenges limit the practical use of these ECVs in helping guide clinicians on treatment and supported the eventual removal of ECVs from the CLSI M100 standards document.

## INTRODUCTION

The *Burkholderia cepacia* complex (BCC) is a group of 24 Gram-negative bacterial species that are best known for their ability to cause lung infections in people with cystic fibrosis (CF) (1-8). BCC also causes opportunistic infections in people without CF and has been linked to outbreaks associated with contaminated medical devices and products (9-16). Treatment of BCC infections is challenging due to antimicrobial resistance, both intrinsic and acquired (17), limited treatment guidance (18), problems with antimicrobial susceptibility testing (AST) methods (19-22), and lack of data correlating AST and clinical outcomes (23). Recently, we and others have shown that AST methods for BCC perform poorly, with disk diffusion (DD), agar dilution (AD), and ETEST failing to correlate with reference broth microdilution (BMD) (19-22). In parallel to these findings, the CF Antimicrobial Resistance (AMR) International Working Group has suggested that AST should not be solely used to treat people with CF because AST results often do not predict clinical outcomes; however, this guidance was extrapolated mostly from data generated with *Pseudomonas aeruginosa* (23, 24). These findings led the Clinical and Laboratory Standards Institute (CLSI) Antimicrobial Susceptibility Testing Subcommittee to remove minimal inhibitory concentration (MIC) and DD breakpoints for BCC from the M100 Performance Standards for Antimicrobial Susceptibility Testing (25). The CLSI decision to remove BCC breakpoints aligns with guidelines from the European Committee on Antimicrobial Susceptibility Testing (EUCAST), which does not recommend AST for BCC nor has published breakpoints for similar reasons (26). However, in the absence of published MIC and DD breakpoints, clinicians may still seek data to help guide antimicrobial therapy for BCC infections.

In the absence of MIC and DD breakpoints, epidemiological cut-off values (ECVs) may be used to help interpret AST results (27-29). CLSI M23 Edition 6 (2023) defines ECVs as MIC or zone diameter values that split microbial populations into those with and without phenotypic resistance mechanisms (non-wild-type or wild-type, respectively); the ECV is the highest MIC or smallest zone diameter value for the wild-type population (30). ECVs may be established for clinically important organisms where there is little pharmacokinetic/pharmacodynamic (PK/PD) and/or clinical outcome data required for establishing breakpoints (27). Indeed, CLSI has established ECVs for some *Candida* and *Aspergillus* species (31). While ECVs do not predict clinical outcomes, they may in some instances be useful to clinicians who are managing difficult-to-treat organisms that lack defined MIC and DD breakpoints. EUCAST has not published ECVs for BCC because the wild-type MIC distribution for antimicrobials used for this species complex is broad (26).

The goal of this study was to establish potential ECVs for BCC against five antimicrobials that had previously published CLSI MIC breakpoints: ceftazidime (CAZ), levofloxacin (LVX), meropenem (MEM), minocycline (MIN), and trimethoprim-sulfamethoxazole (TMP-SMX) (32). These ECVs were published in the CLSI M100 Edition 35 (2025) but were removed from CLSI M100 Edition 36 (2026) for the reasons described here (25, 33).

## MATERIALS AND METHODS

### BCC isolates

MIC data from one thousand eight hundred ninety-six (n=1,896) BCC isolates tested against CAZ, LVX, MEM, MIN, and TMP-SMX were collected from 3 previously published studies and 3 independent laboratories (Table 1). The first two studies (Huse/Jorth *et al*.) (20, 21) included one hundred (n=100) isolates from people with CF (CF isolates) and one hundred five (n=105) isolates from people without CF (non-CF isolates) tested via CLSI reference broth microdilution (34) in triplicate. All isolates were collected from patients in North America (Table 2) and had previously been identified to the species level by whole genome sequencing (Table 1) (20, 21). There were 151 respiratory isolates (100 from people with CF) and 54 non-respiratory isolates (Table 3) (20, 21). In a third study (Wootton *et al*.) (19), 155 CF isolates were tested in triplicate via International Standards Organization BMD (ISO BMD) (19). Isolates were collected in Europe (Table 2) and had previously been identified to the species level via *recA*-based polymerase chain reaction restriction fragment length polymorphism (PCR-RFLP) and matrix-assisted laser desorption/ionization time-of-flight mass spectrometry (MALDI-TOF MS) (BD Bruker MALDI Biotyper) (Table 1) (19). All isolates were collected from sputum (Table 3) (19). In addition to the previously published studies, MIC data were also provided by three other reference laboratories. International Health Management Associates (IHMA; Schaumburg, IL, USA) submitted MIC data for one thousand eleven (n=1,011) isolates that were identified via MALDI-TOF MS (BD Bruker MALDI Biotyper) (Table 1) and tested in single replicate via CLSI reference BMD (34). While the IHMA dataset did not include CF diagnosis, specimen source was included with the data (Table 3). There were six hundred ten (n=610) respiratory isolates, three hundred ninety-nine (n=399) non-respiratory isolates, and two (n=2) isolates from unknown sources (Table 3). Isolates were collected in North America (n=361), Europe (n=258), Latin America (n=169), Asia/West Pacific (n=106), the South Pacific (n=77), Africa (n=29), and the Middle East (n=11) (Table 2). Element Iowa City (JMI Laboratories [JMI]; North Liberty, IA, USA) submitted MIC data for four hundred ninety-nine (n=499) BCC isolates that were identified via MALDI-TOF MS (BD Bruker MALDI Biotyper) (Table 1) and tested in a single replicate via CLSI reference BMD. The isolates were collected at sites in the United States of America (USA, n=340), Europe (n=83), Latin America (n=27), and Asia/West Pacific (n=49) (Table 2). Finally, Microbiologics (St. Cloud, MN, USA) submitted MIC data for twenty-six (n=26) isolates that were collected in the USA (Table 2), identified via MALDI-TOF MS (BD Bruker MALDI Biotyper) (Table 1), and tested via CLSI reference BMD. CF diagnosis and specimen source for the JMI and Microbiologics isolates were not collected. Each study or laboratory contributed different numbers of MIC values depending on the number of isolates submitted and the antimicrobials tested (Table 4).

**Table 1.** *B. cepacia* complex isolates by study or laboratory.

| Identification | Study or Laboratory |  |  |  |  | Total (%) |
| --- | --- | --- | --- | --- | --- | --- |
|  | Huse/Jorth<br>*et al.*<sup>1</sup> | Wootton<br>*et al.*<sup>2</sup> | IHMA | JMI | Microbiologics |  |
| *B. ambifaria* | 0 | 5 | 0 | 0 | 0 | 5<br>(0.3) |
| *B. anthina* | 0 | 3 | 0 | 0 | 0 | 3<br>(0.2) |
| *B. cenocepacia* | 108 | 64 | 0 | 14 | 6 | 192<br>(10.1) |
| *B. cepacia* | 3 | 20 | 0 | 0 | 10 | 33<br>(1.7) |
| *B. cepacia* complex | 0 | 0 | 1011 | 459 | 0 | 1470<br>(77.5) |
| *B. contaminans* | 2 | 2 | 0 | 0 | 0 | 4<br>(0.2) |
| *B. diffusa* | 0 | 1 | 0 | 0 | 0 | 1<br>(0.1) |
| *B. dolosa* | 0 | 4 | 0 | 0 | 0 | 4<br>(0.2) |
| *B. lata* | 0 | 3 | 0 | 0 | 0 | 3<br>(0.2) |
| *B. multivorans* | 92 | 36 | 0 | 26 | 8 | 162<br>(8.5) |
| *B. pyrrocinia* | 0 | 4 | 0 | 0 | 1 | 5<br>(0.3) |
| *B. seminalis* | 0 | 1 | 0 | 0 | 0 | 1<br>(0.1) |
| *B. stabilis* | 0 | 4 | 0 | 0 | 0 | 4<br>(0.2) |
| *B. vietnamiensis* | 0 | 8 | 0 | 0 | 1 | 9<br>(0.5) |
| **Total** | **205** | **155** | **1011** | **499** | **26** | **1896** |
| **(%)** | **(11)** | **(8)** | **(53)** | **(26)** | **(1)** | **(100)** |

1. See references 20 and 21.
2. See reference 19.

**Table 2.** Geographic collection site of *B. cepacia* complex isolates.

| Location | Study or Laboratory |  |  |  |  | Total (%) |
| --- | --- | --- | --- | --- | --- | --- |
|  | Huse/Jorth<br>*et al.* | Wootton<br>*et al.* | IHMA | JMI | Microbiologics |  |
| North America | 205 | 0 | 361 | 340<sup>1</sup> | 26<sup>1</sup> | 932<br>(49) |
| Europe | 0 | 155 | 258 | 83 | 0 | 496<br>(26) |
| Latin America | 0 | 0 | 169 | 27 | 0 | 196<br>(10) |
| Asia/West Pacific | 0 | 0 | 106 | 49 | 0 | 155<br>(8) |
| South Pacific | 0 | 0 | 77 | 0 | 0 | 77<br>(4) |
| Africa | 0 | 0 | 29 | 0 | 0 | 29<br>(2) |
| Middle East | 0 | 0 | 11 | 0 | 0 | 11<br>(1) |
| **Total** | **205** | **155** | **1011** | **499** | **26** | **1896** |

1. Isolates were collected in the USA.

**Table 3.** Specimen sources of *B. cepacia* complex isolates.

| Specimen type | Study or Laboratory |  |  | Total (%) |
| --- | --- | --- | --- | --- |
|  | Huse/Jorth et al. | Wootton et al. | IHMA |  |
| **Respiratory** |  |  |  |  |
| Bronchial or bronchial brushing | 0 | 0 | 136 | 136 (10) |
| Bronchoalveolar lavage | 10 | 0 | 102 | 112 (8.0) |
| Endotracheal aspirate or secretion | 9 | 0 | 190 | 199 (15.0) |
| Endotracheal tube | 4 | 0 | 0 | 4 (0.3) |
| Other | 1 | 0 | 11 | 12 (1.0) |
| Sinus | 1 | 0 | 0 | 1 (0.1) |
| Sputum | 123 | 155 | 169 | 447 (33.0) |
| Throat | 1 | 0 | 0 | 1 (0.1) |
| Tissue | 2 | 0 | 0 | 2 (0.1) |
| Trachea | 0 | 0 | 2 | 2 (0.1) |
| **Total respiratory** | **151** | **155** | **610** | **916 (67.0)** |
| **Non-respiratory** |  |  |  |  |
| Abscess | 3 | 0 | 11 | 14 (1.0) |
| Blood | 24 | 0 | 138 | 162 (12.0) |
| Body fluids | 3 | 0 | 81 | 84 (6.0) |
| Bone | 1 | 0 | 0 | 1 (0.1) |
| Cerebrospinal fluid | 1 | 0 | 0 | 1 (0.1) |
| Dialysate | 1 | 0 | 0 | 1 (0.1) |
| Ear | 2 | 0 | 0 | 2 (0.1) |
| Gastrointestinal | 2 | 0 | 114 | 116 (8.5) |
| Lesion | 0 | 0 | 2 | 2 (0.1) |
| Lymph node | 1 | 0 | 0 | 1 (0.1) |
| Neck drainage | 1 | 0 | 0 | 1 (0.1) |
| Skin, other | 0 | 0 | 3 | 3 (0.2) |
| Urine | 13 | 0 | 38 | 51 (3.7) |
| Wound or ulcer | 2 | 0 | 12 | 14 (1.0) |
| **Total non-respiratory** | **54** | **0** | **399** | **453 (33.0)** |
| **Grand total** | **205** | **155** | **1009** | **1369** |

**Table 4.** Number of minimal inhibitory concentration (MIC) values by study or laboratory and antimicrobial.

| Study or Laboratory | Number MIC Values |  |  |  |  |
| --- | --- | --- | --- | --- | --- |
|  | CAZ (%) | LVX (%) | MEM (%) | MIN (%) | TMP-SMX (%) |
| Huse/Jorth *et al.* | 205<br>(13) | 205<br>(16) | 205<br>(11) | 205<br>(22) | 205<br>(22) |
| Wootton *et al.* | 155<br>(10) | 0<br>(0) | 155<br>(9) | 155<br>(17) | 155<br>(16) |
| IHMA | 642<br>(42) | 534<br>(42) | 901<br>(50) | 62<br>(7) | 85<br>(9) |
| JMI | 498<br>(33) | 499<br>(39) | 499<br>(28) | 491<br>(54) | 498<br>(53) |
| Microbiologics | 26<br>(2) | 26<br>(2) | 26<br>(1) | 0<br>(0) | 0<br>(0) |
| **Total (%)** | **1526**<br>**(100)** | **1264**<br>**(100)** | **1786**<br>**(100)** | **913**<br>**(100)** | **943**<br>**(100)** |

Abbreviations: MIC=minimal inhibitory concentration; CAZ=ceftazidime; LVX=levofloxacin; MEM=meropenem; MIN=minocycline; TMP-SMX=trimethoprim-sulfamethoxazole. Red font indicates antimicrobials where $\geq 50\%$ of the MICs were contributed by one laboratory.

### Determination of ECVs

Three analyses were used to determine BCC ECVs for CAZ, LVX, MEM, MIN, and TMP-SMX. CLSI M23 Edition 6 (2023) guidelines were followed with exceptions or limitations as noted (see Results and Discussion) (30). For each analysis, ECVs were determined using the iterative statistical method with ECOFFinder v2.1^35^ (https://clsi.org/meetings/susceptibility-testing-subcommittees/ecoffinder/). As noted in CLSI M23 Edition 6 (2023), a 97.5% threshold was applied to determine ECVs (30).

In the first analysis, MIC data from all isolates listed in Table 1 were pooled. For the Huse/Jorth *et al*. (20, 21) and Wootton *et al*. (19) data, the mode MIC value from triplicate testing was used, while for the IHMA, JMI, and Microbiologics data, the single replicate value was used. In this analysis, data were not weighted. In the second analysis, the same analysis was performed as above, except data were weighted for MEM, MIN, and TMP-SMX because one laboratory contributed ≥ 50% of the MIC data. Susceptibility data were weighted by calculating the percentage of isolates at each MIC for each contributing study or laboratory, multiplied by one hundred, and rounded to the nearest whole number. In the third analysis, JMI data for TMP-SMX were removed from the pooled dataset because MIC data were truncated. ECVs were then determined without JMI data for TMP-SMX.

## RESULTS

### BCC isolates

MIC data for 1,896 BCC isolates were collected from 3 previously published studies and 3 reference laboratories (Table 1). IHMA contributed the most data (n=1,011, 53%), followed by JMI (n=499, 26%), Huse/Jorth *et al*. (n=205, 11%) (20, 21), Wootton *et al*. (19) (n=155, 8%), and Microbiologics (n=26, 1%). Most isolates were identified to the complex level (n=1,470; 77.5%), while the remaining isolates (n=426; 22.5%) were identified to the species level (Table 1). Of the isolates that were identified to the species level, *B. cenocepacia* (n=192, 10.1%), *B. multivorans* (n=162, 8.5%), and *B. cepacia* (n=33, 1.7%) were the most common (Table 1). Thirty-nine (n=39; 2%) other isolates were identified to the species level: *B. ambifaria*, n=5; *B. anthina*, n=3; *B. contaminans*, n=4, *B. diffusa*, n=1; *B. dolosa*, n=4, *B. lata*, n=3; *B. pyrrocinia*, n=5; *B. seminalis*, n=1; *B. stabilis*, n=4, and *B. vietnamiensis*, n=9 (Table 1). Isolates were collected from seven geographic locations, with most isolated in North America (n=932, 49%), followed by Europe (n=496, 26%), Latin America (n=196, 10%), Asia/West Pacific (n=155, 8%), South Pacific (n=77, 4%), Africa (n=29, 2%), and the Middle East (n=11, 1%) (Table 2). For most isolates, the specimen source was identified (n=1,369, 72%), with the majority collected from respiratory sources (n=916, 67%) (Table 3). Sputum was the most common respiratory source (n=447, 33%), followed by endotracheal aspirate or secretion (n=199, 15%) (Table 3). Of the non-respiratory specimens (n=453, 33%), most were from blood (n=162, 12%), followed by gastrointestinal sources (n=116, 8.5%) (Table 3). For most isolates, it was unknown whether isolates were collected from a person with CF (n=1641, 87%). The number of MIC values for each antimicrobial varied depending on the study or laboratory (Table 4). For CAZ, most data were provided by IHMA (n=642, 42%), followed by JMI (n=498, 33%), Huse/Jorth *et al*. (20, 21) (n=205, 13%), Wootton *et al*. (19) (n=155, 10%), and Microbiologics (n=26, 2%). Similarly, for LVX, IHMA (n=534, 42%) and JMI (n=499, 39%) provided the most data, followed by Huse/Jorth *et al*. (20, 21) (n=205, 16%) and Microbiologics (n=26, 2%). IHMA contributed the most data (n=901, 50%) for MEM, followed by JMI (n=499, 28%), Huse/Jorth *et al*. (20, 21) (n=205, 11%), Wootton *et al*. (19) (n=155, 9%), and Microbiologics (n=26, 1%). For MIN, JMI contributed the most data (n=491, 54%), followed by Huse/Jorth *et al*. (20, 21) (n=205, 22%), Wootton *et al*. (19) (n=155, 17%), and IHMA (n=62, 7%). Finally, for TMP-SMX, JMI contributed the most data (n=498, 53%), followed by Huse/Jorth *et al*. (20, 21) (n=205, 22%), Wootton *et al*. (19) (n=155, 16%), and IHMA (n=85, 9%).

### Wild-type distributions and determination of ECVs

MIC distributions were produced for each antimicrobial using the mode MIC from Huse/Jorth *et al*. (20, 21) and Wootton *et al*. (19) or single replicate MIC from IHMA, JMI, and Microbiologics (Figure 1). While the MIC distribution for MEM followed a log normal distribution, the MIC distributions for CAZ, LVX, MIN, and TMP-SMX showed upper end tailing due to more isolates with higher MICs (Figure 1). If the now obselete MIC breakpoints were applied, a large proportion of isolates were not susceptible (intermediate or resistant), with 26%, 40%, 28%, 27%, and 28% not susceptible to CAZ, LVX, MEM, MIN, and TMP-SMX, respectively.

**Figure 1.**
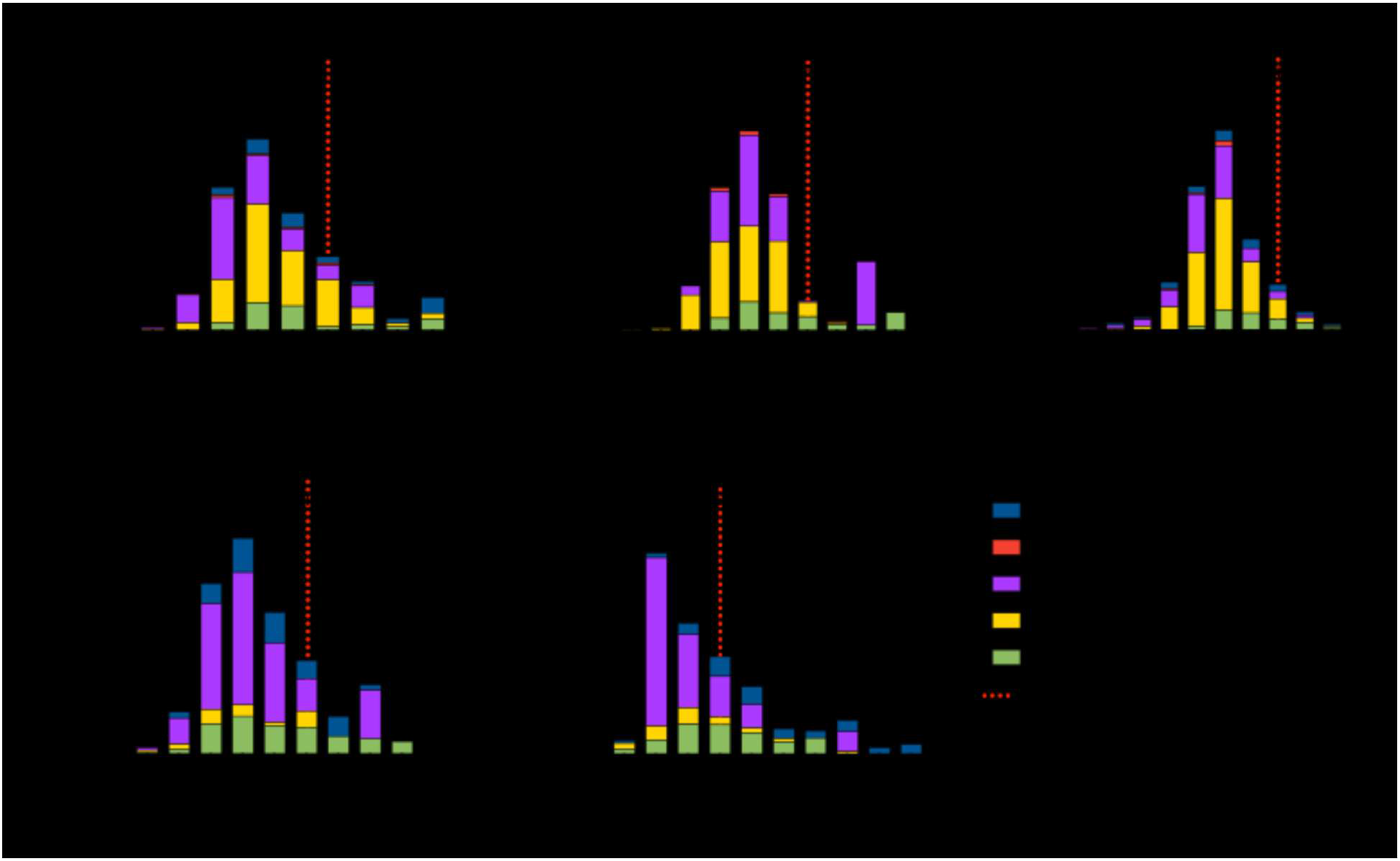
Broth microdilution (BMD) minimal inhibitory concentration (MIC) distributions for *B. cepacia* complex (BCC) isolates for CAZ, LVX, MEM, MIN, and TMP-SMX. Histograms show BMD MIC distributions for ceftazidime (CAZ), levofloxacin (LVX), meropenem (MEM), minocycline (MIN), and trimethoprim-sulfamethoxazole (TMP-SMX). The total number of isolates analyzed for each antibiotic are indicated on each graph (n=number). Black dashed lines indicate obsolete BCC MIC breakpoints from CLSI M100 ED34:2024 (32). Red dashed lines indicate the ECV for non-weighted data using a 97.5% threshold. Wootton *et al*. (19) study (dark blue); Microbiologics (red); JMI: JMI Laboratories (purple); IHMA: International Health Management Associates (yellow); Huse/Jorth *et al*. (20, 21) study (green); S=susceptible; I=intermediate; R=resistant.

For MEM, MIN, and TMP-SMX, ≥ 50% of the data points were generated from one laboratory. In these cases, CLSI M23 Edition 6 (2023) guidelines state that weighting the data should be considered before pooling (30). To determine whether weighting data impacted determination of ECVs, analyses were performed with and without data weighting for MEM, MIN, and TMP-SMX. Per CLSI M23 Edition 6 (2023) guidelines, a 97.5% threshold was applied to determine ECVs (30). For the non-weighted data, the ECVs were 16 μg/ml for CAZ, 8 μg/ml for LVX, 16 μg/ml for MEM, 8 μg/ml for MIN, and 2 μg/ml for TMP-SMX (Figure 1 and Table 5). These ECVs would have fallen into the following categories if previously published CLSI MIC breakpoints were applied: CAZ 16 μg/ml, intermediate; LVX, 8 μg/ml, resistant; MEM 16 μg/ml, resistant; MIN, 8 μg/ml, intermediate, and TMP-SMX, 2 μg/ml, susceptible. Note that these were the ECVs published in M100 Edition 35 (2025) (25). When data were weighted, the ECVs for MEM and MIN were the same (16 μg/ml and 8 μg/ml, respectively), but the ECV for TMP-SMX increased from 2 μg/ml to 8 μg/ml (Figure 2 and Table 5).

**Table 5.** Epidemiological cut-off values (ECV) by analysis.

| Antimicrobial | Non-weighted ECV<br>(µg/mL)<sup>1</sup> | Weighted ECV<br>(µg/mL) | ECV, JMI data removed<br>(µg/mL) |
| --- | --- | --- | --- |
| CAZ | 16 | NA | NA |
| LVX | 8 | NA | NA |
| MEM | 16 | 16 | NA |
| MIN | 8 | 8 | NA |
| TMP-SMX | 2 | 8 | 16 |

1. Published in CLSI M100 Edition 35 (2025)<sup>25</sup>.
Abbreviations: ECV=epidemiological cut-off value. CAZ=ceftazidime; LVX=levofloxacin; MEM=meropenem; MIN=minocycline; TMP-SMX=trimethoprim-sulfamethoxazole. NA=not applicable.

**Figure 2.**
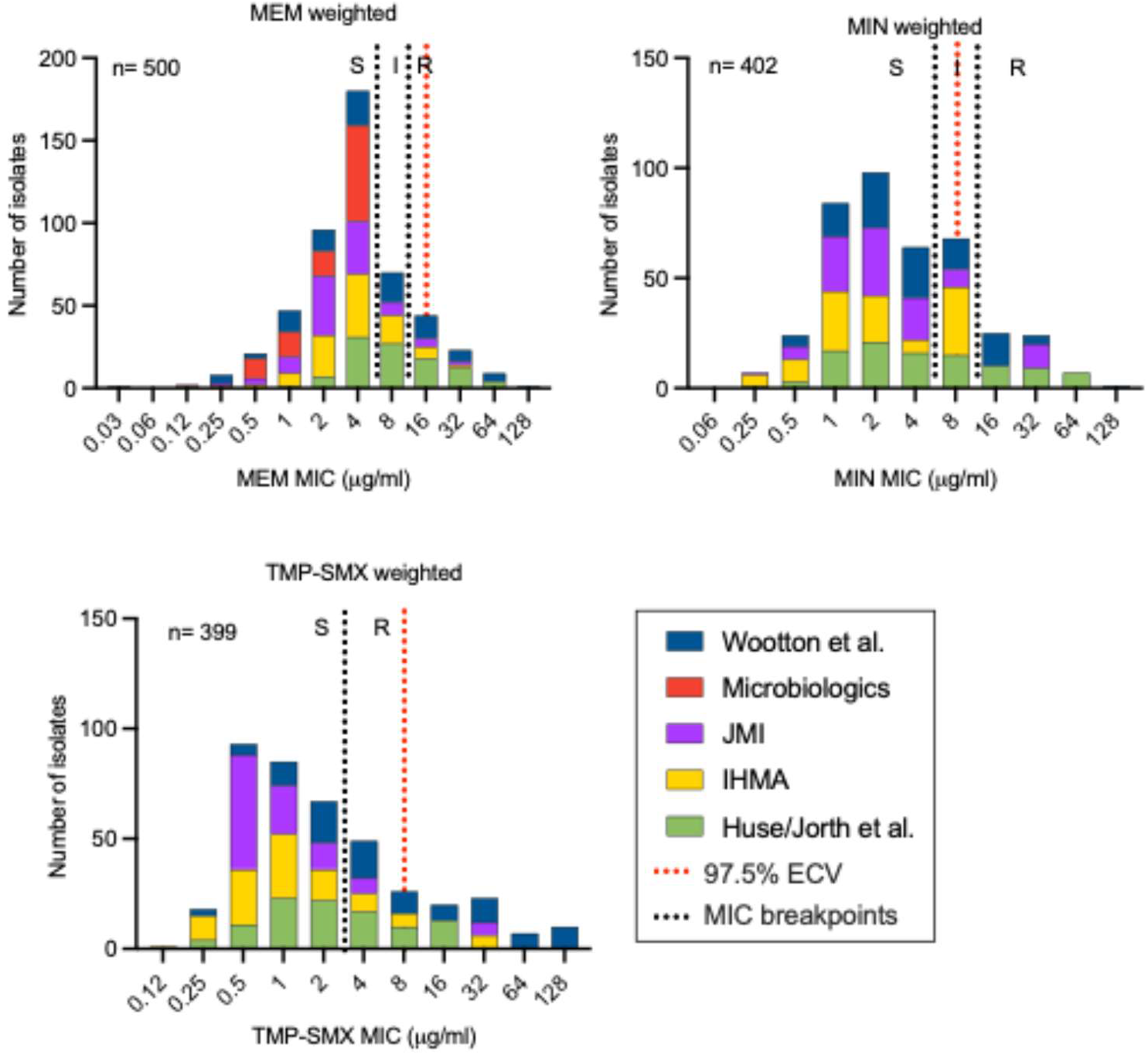
Determination of epidemiological cutoff values (ECVs) for meropenem (MEM), minocycline (MIN), and trimethoprim-sulfamethoxazole (TMP-SMX) after weighting pooled minimal inhibitory concentration (MIC) data. Histograms show weighted MIC distributions for meropenem (MEM), minocycline (MIN), and trimethoprim-sulfamethoxazole (TMP-SMX) (see Materials and Methods). The total number of isolates after for each antibiotic after weighting are indicated on each graph (n=number). Black dashed lines indicate obsolete *B. cepacia* complex MIC breakpoints from CLSI M100 ED34:2024 (32). Red dashed lines indicate the ECV for weighted data using a 97.5% threshold. Wootton *et al*. (19) study (dark blue); Microbiologics (red); JMI: JMI Laboratories (purple); IHMA: International Health Management Associates (yellow); Huse/Jorth *et al*. (20, 21) study (green); S=susceptible; I=intermediate; R=resistant.

When further analyzing MIC data for TMP-SMX, it was noted that the JMI data were truncated on the lower end of the MIC distribution, which may impact the ECV. Therefore, a third analysis was performed to determine the TMP-SMX ECV where the JMI data (n=498) were removed from the pooled dataset. The ECV was then determined (Figure 3 and Table 5). When the JMI data were removed, the ECV for TMP-SMX was 16 μg/ml (Figure 3 and Table 5).

**Figure 3.**
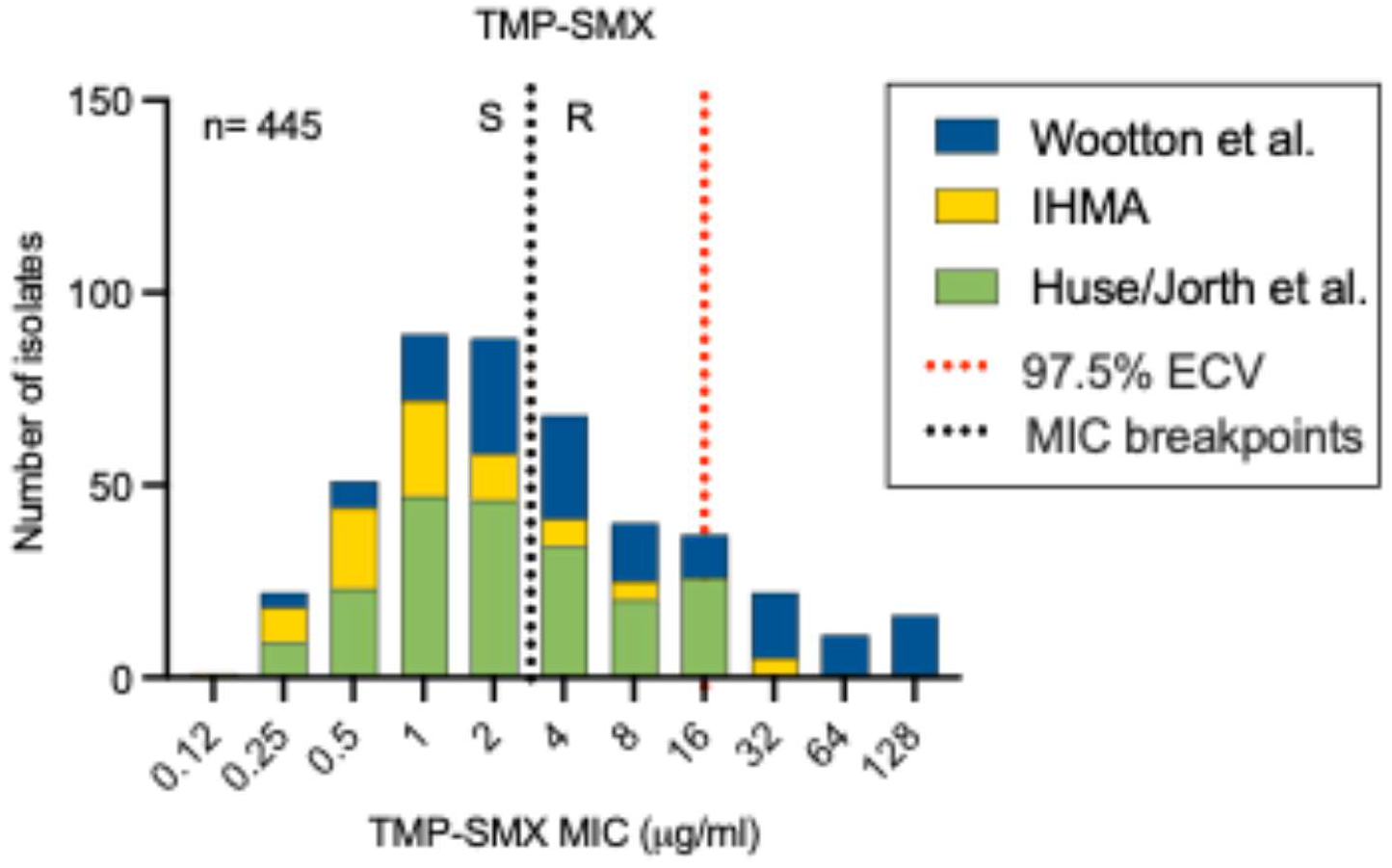
Determination of epidemiological cutoff values (ECVs) for TMP-SMX after removing truncated JMI data from pooled minimal inhibitory concentration (MIC) data. Histograms show MIC distributions for trimethoprim-sulfamethoxazole (TMP-SMX) after removal of truncated JMI data from the pooled MIC dataset (see Materials and Methods). The total number of isolates after omitting JMI data is indicated (n=number). Black dashed lines indicate obsolete *B. cepacia* complex MIC breakpoints from CLSI M100 ED34:2024 (32). Red dashed lines indicate the ECV using a 97.5% threshold. Wootton *et al*. (19) study (dark blue); IHMA: International Health Management Associates (yellow); Huse/Jorth *et al*. (20, 21) study (green); n=number; S=susceptible; I=intermediate; R=resistant.

## DISCUSSION

Since 2024, CLSI AST guidelines for BCC have undergone significant changes. Based on data showing that DD does not correlate with CLSI reference BMD, DD breakpoints were removed from CLSI M100 Edition 34 (2024) (32). Similarly, when studies showed that reference AD and BMD did not correlate, MIC breakpoints were removed from CLSI M100 Edition 35 (2025) (25). The ECVs established here were added to CLSI M100 Edition 35 (2025) (25) but were removed from CLSI M100 Edition 36 (2026) (33), as discussed below.

CLSI M23 Edition 6 (2023) recommends that data for establishing ECVs should include the following: a minimum of 100 isolates collected from 3 independent sites identified to the species level and tested by CLSI reference methods (30). In this study, we were able to collect MIC data from 1,896 BCC isolates tested previously at 5 sites. This number was well above the minimum requirement and exceeded previous ECV analyses for *Aspergillus* species (n=22-833 depending on species) (36) but was far less than other ECV analyses for *Candida* species (n=11,241-15,269) (37, 38). Isolates were collected from seven different geographic regions and included a wide variety of specimen types, which increased robustness of the dataset. Notably, the MIC distributions for CAZ, LVX, MIN, and TMP-SMX were and included a high number of not susceptible isolates. While it is expected to find not susceptible isolates in people with CF (17), only 13% of isolates studied here were identified as isolates from people with CF; it was unknown whether the remaining 87% of isolates were collected from a person with CF. Next, only 22.5% of isolates were identified to the species level. Therefore, ECVs could only be established for the *B. cepacia* complex and may be biased toward more highly represented species like *B. cenocepacia, B. multivorans*, and *B. cepacia* (20% of isolates identified to the species level). Given that these species are the most frequently isolated, this limitation may not be greatly problematic but does not follow CLSI M23 Edition 6 (2023) guidelines, which states that isolates should be identified to the species level (30, 39, 40). Finally, the isolates from the Wootton *et al*. study (19) were tested by ISO BMD, which differs in very minor ways from the CLSI reference BMD method and was therefore not expected to yield different results. We included these isolates to increase the number of MICs in the pooled dataset and because they were identified to the species level. Overall, our dataset was constrained by the infrequent isolation of these organisms compared to other non-fermenting Gram-negative rods like *P. aeruginosa* and laboratory variability in identification to the species level (41-45).

Because DD and MIC breakpoints were removed from the CLSI M100 (25, 32), there was significant concern that clinicians would have no guidance for these often difficult-to-treat infections. Therefore, ECVs for CAZ (16 μg/ml), LVX (8 μg/ml), MEM (16 μg/ml, not weighted), MIN (8 μg/ml, not weighted), and TMP-SMX (2 μg/ml, not weighted) were included in CLSI M100 Edition 35 (2025) (25) despite the limitations described above. Additionally, the published ECVs for CAZ, LVX, MEM, and MIN were higher than the obsolete susceptible breakpoints (CAZ, 8 μg/ml; LVX 2 μg/ml; MEM, 4 μg/ml; and MIN 4 μg/mL). In such cases, if the ECV were to be applied clinically for patient care, some of the wild type population would fall within the resistant category, potentially leading to misguided therapy. Footnotes were included in the CLSI M100 Edition 35 (2025) to address these limitations (25).

Pharmacokinetic/pharmacodynamic (PK/PD) data further underscore the limited clinical applicability of these ECVs. Although BCC-specific data are limited, studies in other non-fermenting Gram-negative organisms provide relevant context. Against *Stenotrophomonas maltophilia*, the probability of target attainment (PTA) for TMP-SMX declines at MICs ≥ 1 mg/L and exposures required to achieve a 1-log10 CFU/mL reduction were not quantifiable (46). Similarly, for LVX against *S. maltophilia*, PTA for achieving a 1-log_10_ kill at an MIC of 2 mg/L was 26.6% (47). Comparable findings are reported for MEM, with PTA declining at MICs > 2 mg/L in *P. aeruginosa* (48). High-dose MIN is predicted to provide suboptimal PK/PD target attainment for MICs > 1 mg/L against Gram-negative bacteria (49). For CAZ, exposures sufficient for an MIC of 16 mg/L may be achieved with high-dose, prolonged infusion, but not consistently across patient populations (50). Together, these data indicate that PK/PD target attainment is often not achievable at MIC values below or near the proposed BCC ECVs.

We performed additional analyses for BCC ECVs to address the limitations described above. Data weighting did not impact the MEM or MIN ECVs (Table 5). However, data weighting increased the TMP-SMX ECV from 2 μg/mL to 8 μg/mL, and removal of the truncated JMI data increased the ECV to 16 μg/ml (Table 5). ECVs could be confidently established for CAZ, LVX, MEM, and MIN at 16 μg/ml, 8 μg/ml, 16 μg/ml, and 8 μg/ml, respectively, but variation in TMP-SMX ECVs by analysis (2-16 μg/mL) limited confidence in its interpretation. When the ECVs were re-visited by the CLSI AST subcommittee, there were concerns about 1) the high values of the ECVs and whether they would be clinically achievable targets (46-51); 2) variation of the TMP-SMX ECV depending on analysis; and 3) if clinicians would use ECVs as breakpoints. It was decided that ECVs should be removed from CLSI M100 Edition 36 (2026) due to these concerns (33, 52). Overall, significant limitations within the BCC ECV dataset, elevated MIC distributions for these organisms, and the lack of clinically achievable ECVs substantially limit their practical utility, and thus the ECVs were removed from the CLSI M100 standards documuent. Given these limitations, interpretation of AST results for these organisms remains complex, and multidisciplinary input from clinical microbiology, antimicrobial stewardship, and infectious diseases is advised.

## Acknowledgements

This research was supported by grants to PJ and HH from the Cystic Fibrosis Foundation (JORTH19I0, JORTH23XX0, and NAREN24R20), JJL is supported by grants from the Cystic Fibrosis Foundation, and ACMG was supported by a fellowship from the Cystic Fibrosis Foundation (MILESI21F0). We thank the CLSI Ad Hoc Working Group on BCC AST for their helpful feedback during the study. We thank the Canadian *Burkholderia cepacia* Research and Referral Repository (funded by Cystic Fibrosis Canada). We also thank the people with cystic fibrosis who contributed bacterial isolates to the Cystic Fibrosis Foundation *Burkholderia cepacia* Research Laboratory and Repository at the University of Michigan.

